# Modeling missing cases and transmission links in networks of extensively drug-resistant tuberculosis in KwaZulu-Natal, South Africa

**DOI:** 10.1101/655969

**Authors:** Kristin N. Nelson, Neel R. Gandhi, Barun Mathema, Benjamin A. Lopman, James C.M. Brust, Sara C. Auld, Nazir Ismail, Shaheed Vally Omar, Tyler S. Brown, Salim Allana, Angie Campbell, Pravi Moodley, Koleka Mlisana, N. Sarita Shah, Samuel M. Jenness

## Abstract

The transmission patterns of drug-resistant tuberculosis (TB) remain poorly understood, despite over half a million incident cases in 2017. Modeling TB transmission networks can provide insight into the nature and drivers of transmission, but incomplete and non-random sampling of TB cases can pose challenges to making inferences from epidemiologic and molecular data. We conducted a quantitative bias analysis to assess the effect of missing cases on a transmission network inferred from *Mtb* sequencing data on extensively drug-resistant (XDR) TB cases in South Africa. We tested scenarios in which cases were missing at random, differentially by clinical characteristics, or differentially by transmission (i.e., cases with many links were under or over-sampled). Under the assumption cases were missing at random, cases in the complete, modeled network would have had a mean of 20 or more transmission links, which is far higher than expected, in order to reproduce the observed, partial network. Instead, we found that the most likely scenario involved undersampling of high-transmitting cases, and further models provided evidence for superspreading behavior. This is, to our knowledge, the first study to define and assess the support for different mechanisms of missingness in a study of TB transmission. Our findings should caution interpretation of results of future studies of TB transmission in high-incidence settings, given the potential for biased sampling, and should motivate further research aimed at identifying the specific host, pathogen, or environmental factors contributing to superspreading.

---

Tuberculosis (TB) is the leading infectious cause of death worldwide.(1) The ongoing transmission of extensively drug-resistant (XDR) tuberculosis, which is resistant to both first and second-line antibiotics, represents a severe threat to public health. South Africa has among the highest rates of TB and HIV globally and KwaZulu-Natal province has the highest XDR TB burden in South Africa (3 per 100,000). (2–5) In South Africa and elsewhere, the majority of drug-resistant TB cases are due to transmission of already-resistant strains, rather than inadequate treatment.(6, 7) This underscores the importance of locating where TB transmission occurs in order to develop interventions, including environmental control measures, targeting key transmission locations and at-risk groups.(8) These efforts will involve identifying both sites of transmission and attributes of individuals who are most likely to transmit to their susceptible contacts.

Identifying discrete transmission events is a challenge given the airborne transmission route of TB. However, bacterial whole genome sequencing allows for characterization of differences between *Mycobacterium tuberculosis* (*Mtb*) sequences at the level of individual base pairs. Cases with similar *Mtb* sequences are likely to be linked through transmission, and collectively, such links can be used to infer networks of transmission events.(9) Many previous studies have inferred transmission events using social contact or molecular sequence data, but a key limitation of these studies in high-incidence settings is that it is practically impossible to identify all TB cases. Half of TB cases are estimated to be undiagnosed, and this proportion is likely higher for XDR TB given the need for culture-based drug susceptibility testing that is not universally available(10–12). Further, epidemiologic or sequencing information may be missing, either because a clinical sample could not be collected or because a case died prior to diagnosis or study enrollment (a particular concern with XDR TB, given its 28% survival rate)(13)). Thus, a major challenge of characterizing TB transmission networks in these settings is inferring a complete, or at least representative, set of transmission links from incomplete data. If the network constructed from incomplete data poorly resembles the true transmission network, inferences about transmission patterns may be biased.

A modeling approach to the problem of missing network data could provide insight into what the structure of the complete transmission network, had it been measured, would have looked like. Missing network data is different from missing data in traditional epidemiologic studies, because the dependence among cases in a network violates the assumption that data are independent and identically distributed. As a result, even if cases are missing completely at random from the network, inference from a partially sampled transmission network could be biased. However, inference may still be made about the complete network if certain conditions are met.(14) Most importantly, sampled cases must not differ systematically from unsampled cases with respect to their transmission potential.(14) This may occur if, for example, undiagnosed cases have longer infectious periods and therefore contribute disproportionately to transmission as compared to diagnosed cases who receive prompt treatment. Failing to sample these highly connected cases could have a pronounced effect on network structure and, as a result, bias conclusions made from the empirical network.(15) However, bias may be mitigated if it can be quantified, by identifying characteristics of cases that are undersampled and using this information to provide insight into the structure of the true network. For example, TB cases without detectable mycobacteria in sputum (‘smear-negative’) are both less infectious and more difficult to diagnose than smear-positive TB cases, leading to scenario in which diagnosed cases may be responsible for more transmission than undiagnosed and unsampled cases.(16) Understanding characteristics of cases that may be missing is a first step in evaluating the robustness of empirical network data on TB cases in different settings. Lastly, understanding the composition of complete transmission networks allows us to test hypotheses about drivers of TB transmission. Superspreading in TB is increasingly recognized as an important phenomenon shaping transmission dynamics, though it is difficult to capture definitive evidence of superspreading events.(17, 18) Detecting signatures of this phenomenon in transmission networks will improve our understanding of its role in driving TB transmission.

In this study, we used data from the <u>T</u>ransmission study of <u>X</u>DR TB (TRAX), which enrolled culture-confirmed XDR TB cases diagnosed from 2011 to 2014 in KwaZulu-Natal, South Africa. We constructed an empirical transmission network based on *Mtb* sequence data, then used network models to infer ‘complete’ transmission networks based on different assumptions about how data was missing from the empirical network. We also tested several models including a ‘superspreading’ factor to understand its impact on network structure. Our goal was to identify the typology of missingness most consistent with the empirical network, to assess whether it is likely to be representative of true XDR TB transmission patterns.

## METHODS

### Study design and procedures

Detailed methods of TRAX procedures have been published.(6, 19) Briefly, we identified 1027 XDR TB cases through the single referral laboratory that conducts drug-susceptibility testing (DST) for all public healthcare facilities in the province and selected a convenience sample of 404 cases. All participants provided written informed consent; for deceased or severely ill participants, consent was obtained from next-of-kin. We interviewed participants and performed medical record review to collect demographic and clinical information. The diagnostic XDR TB isolate was obtained for all enrolled participants. Raw paired-end sequencing reads were generated on the Illumina (MiSeq) platform and aligned to the H37Rv reference genome (NC_000962.3). Single nucleotide polymorphisms (SNPs) were detected using standard pairwise resequencing techniques (Samtools v0.1.19) against the reference.(20) 344 cases had *Mtb* sequences that passed all quality filters.

### Constructing the empirical network using Mtb sequence data

We defined a genomic link as a pair of XDR TB cases with 5 or fewer SNP differences between their *Mtb* sequences. We constructed genomic transmission networks of TRAX cases, in which each *node* in the network represents a case and each *edge* represents a potential transmission event (sum of the source and forward transmission links). The degree of each node is the number of edges per case; the degree distribution represents the edge count across all nodes in the network. We conceptualized this empirical genomic network as a subset of the cases and links in the true, complete transmission network that includes all XDR TB cases and transmission events in KwaZulu-Natal during the study period.

### Exponential random graph models (ERGMs)

Conventional statistical models assume that the characteristics of each subject are independent from others’, an assumption that is not met in disease transmission networks. In a transmission network, the unit of interest is a transmission link, which consists of two cases whose attributes may be correlated. ERGMs are a tool for statistically modeling the propensity of links to form between nodes (cases) in a network, accounting for the inherent correlation among attributes of cases within the network. We used ERGMs to express the probability that a transmission link occurs between two cases in the network as a function of demographic and clinical characteristics of each case. This approach allowed us to understand the structure of the complete network as a joint function of the empirical data and assumptions about missingness.

We used infectiousness estimates from the literature to define target statistics for the degree of a case based on their attributes (e.g., smear-negative cases had, on average, 25% fewer edges as compared to smear-positive cases).(12, 21) If there was limited information in the literature for a given attribute (cough duration, strain type), we used data from TRAX to define target statistics (Technical Appendix). We specified models under each missing data scenario. Since the mean degree (number of links per case) in the complete network was unknown, we tested each scenario across a range of mean degrees. From each scenario, we simulated 1,000 complete transmission networks (Figure 1). ERGMs were constructed using the *ergm* R package.(22, 23) Code is available at https://github.com/kbratnelson/tb-ergms.

**Figure 1.**
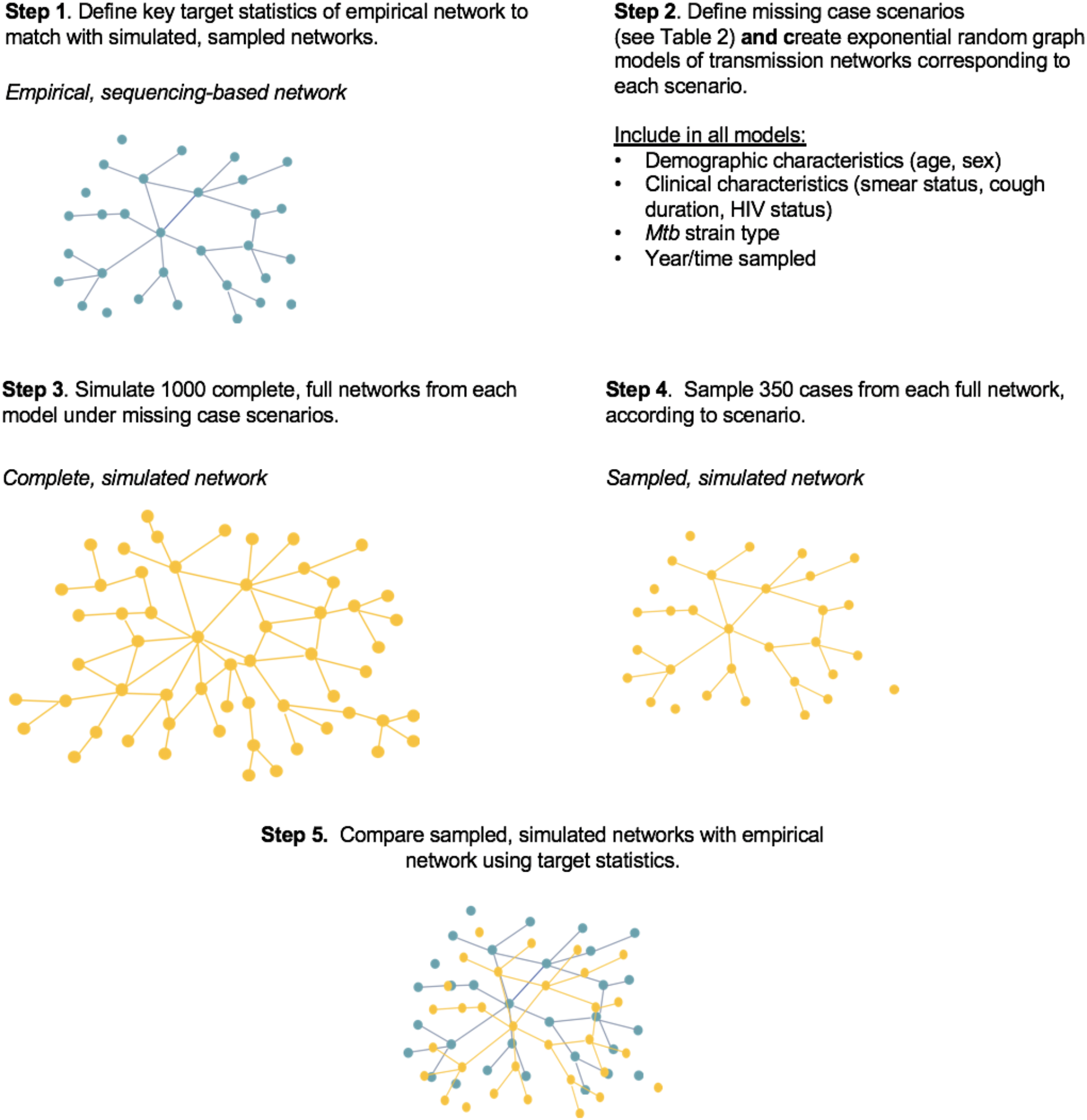
Schematic Representation of Modeling and Sampling Methods

To model complete transmission networks, we needed to estimate the total number of diagnosed and undiagnosed XDR TB cases in KwaZulu-Natal during the study period (2011–2014). We used data from the South African Tuberculosis Drug Resistance Survey to estimate the number of diagnosed XDR TB cases and active case-finding studies to estimate the number of additional, undiagnosed cases (Technical Appendix).(24) We assumed a complete transmission network size of 2000 cases for our primary analyses.

### Missing data scenarios

We modeled three different scenarios under which cases may be missing from the empirical network. First, we assumed that cases were missing at random (Scenario 1, Table 1). Second, we assumed systematic oversampling of cases either involved in many transmission events (‘high-transmitters’) or few transmission events (‘low-transmitters’) (Scenario 2).

**Table 1.** Missing Case Scenarios.

| Scenario | Changes to complete transmission network model or sampling procedures |
| --- | --- |
| 1. Cases missing at random | No changes to model terms. |
| 2. Cases missing by transmission | No changes to model terms. Sample from complete network nonrandomly using degree to define sampling weights.<br>**a.** Highly connected cases ('high-transmitters') more likely to be sampled: sampling weighted by degree<br>**b.** Poorly connected cases ('low-transmitters') more likely to be sampled: sampling weighted by inverse degree |
| 3. Cases missing by smear status | No changes to model terms. Increase proportion of smear-positive cases in complete network relative to sampled network. |
| 4. Latent, unmeasured (superspreading) factor | Add model term corresponding to unmeasured factor strongly related to transmission in a minority of cases. Vary strength and prevalence of factor.<br>**a.** Unmeasured factor that increases transmission by factor of 10 (prevalence: 10%, 20%, 30%)<br>**b.** Unmeasured factor that increases transmission by factor of 20 (prevalence: 10%, 20%, 30%)<br>**c.** Unmeasured factor that increases transmission by factor of 40 (prevalence: 10%) |

Third, we hypothesized that cases were sampled differentially based on smear status (Scenario 3). Smear-negative cases are more difficult to diagnose and may therefore be underrepresented in empirical transmission networks.(21)

The final scenario did not make assumptions about case sampling, but modeled a latent, unmeasured factor strongly related to the likelihood of transmission (Scenario 4). This tested the hypothesis that an unmeasured characteristic in a minority of cases, representing superspreading, could explain the structure of the empirical transmission network.

### Sampling from Modeled Networks

From each modeled, complete network, we sampled a similar number of cases (350) as in our empirical network. We aimed to determine which scenario produced sampled networks that most closely matched the empirical network with respect to the degree distribution. To compare the empirical network with the simulated and sampled networks, we compared locations of the quantiles (10%, 25%, 50%, 100%) of the degree distribution.

### Additional Sensitivity Analyses

Because there is considerable uncertainty about the genomic threshold that should be used to define a direct TB transmission event, we conducted the same analysis using a more stringent SNP threshold (3 SNPs). We also tested the sensitivity of our results to assumptions about the size of the complete transmission network, which is unknown, by decreasing (n = 1500) and increasing (n = 4000) complete network size.

## RESULTS

The empirical genomic network comprised 344 TRAX cases with 1084 genomic links, or edges. Each case had an average of 6.3 edges (the overall network mean degree), and 182 (53%) cases in the network had at least one edge. The 25th percentile of the degree distribution was located at 0, the 50^th^ percentile (median) was at 1, the 75^th^ percentile at 7 (Table 2). The most highly linked case had a degree of 62; and 62 (18%) cases had 10 or more links.

**Table 2.**
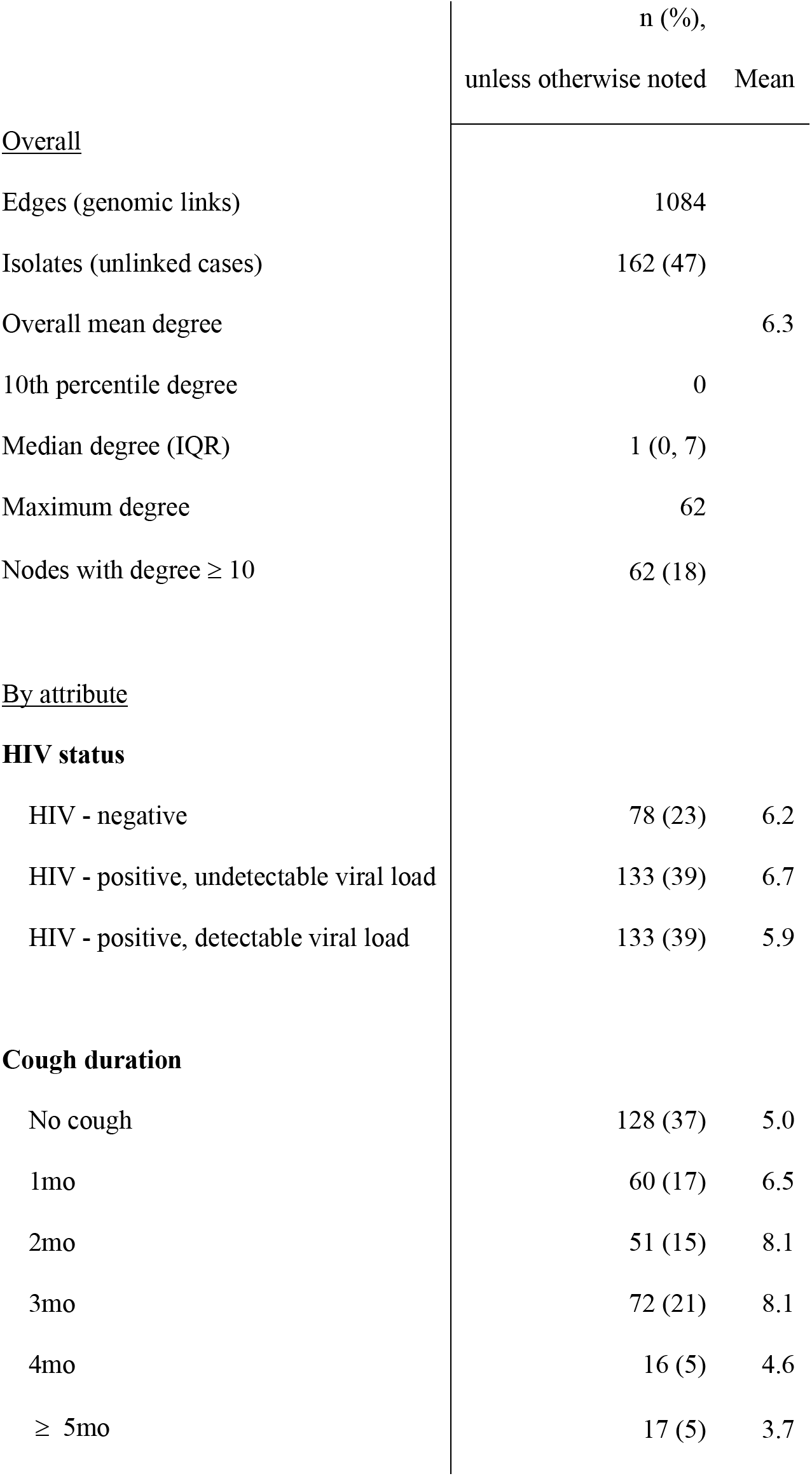
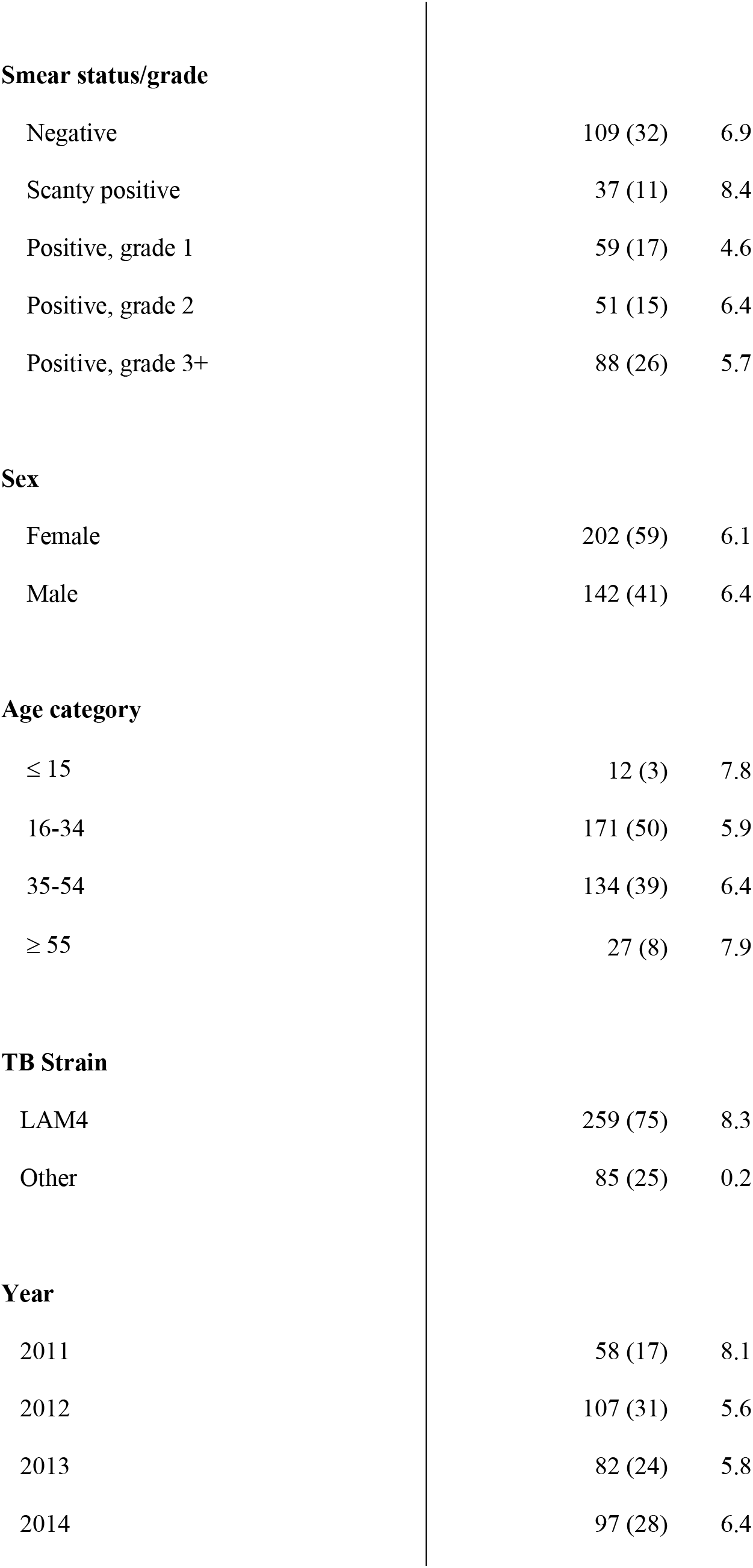
Descriptive Characteristics of the Empirical Sequencing-Based Network of XDR TB cases from the TRAX study in KwaZulu-Natal, South Africa, 2011-2016.

|  | n (%),<br>unless otherwise noted | Mean |
| --- | --- | --- |
| <u>Overall</u> |  |  |
| Edges (genomic links) | 1084 |  |
| Isolates (unlinked cases) | 162 (47) |  |
| Overall mean degree |  | 6.3 |
| 10th percentile degree | 0 |  |
| Median degree (IQR) | 1 (0, 7) |  |
| Maximum degree | 62 |  |
| Nodes with degree $\geq 10$ | 62 (18) | |
| <u>By attribute</u> |  |  |
| **HIV status** |  |  |
| HIV - negative | 78 (23) | 6.2 |
| HIV - positive, undetectable viral load | 133 (39) | 6.7 |
| HIV - positive, detectable viral load | 133 (39) | 5.9 |
| **Cough duration** |  |  |
| No cough | 128 (37) | 5.0 |
| 1mo | 60 (17) | 6.5 |
| 2mo | 51 (15) | 8.1 |
| 3mo | 72 (21) | 8.1 |
| 4mo | 16 (5) | 4.6 |
| $\geq 5$mo | 17 (5) | 3.7 |
| **Smear status/grade** |  |  |
| Negative | 109 (32) | 6.9 |
| Scanty positive | 37 (11) | 8.4 |
| Positive, grade 1 | 59 (17) | 4.6 |
| Positive, grade 2 | 51 (15) | 6.4 |
| Positive, grade 3+ | 88 (26) | 5.7 |
| **Sex** |  |  |
| Female | 202 (59) | 6.1 |
| Male | 142 (41) | 6.4 |
| **Age category** |  |  |
| ≤ 15 | 12 (3) | 7.8 |
| 16-34 | 171 (50) | 5.9 |
| 35-54 | 134 (39) | 6.4 |
| ≥ 55 | 27 (8) | 7.9 |
| **TB Strain** |  |  |
| LAM4 | 259 (75) | 8.3 |
| Other | 85 (25) | 0.2 |
| **Year** |  |  |
| 2011 | 58 (17) | 8.1 |
| 2012 | 107 (31) | 5.6 |
| 2013 | 82 (24) | 5.8 |
| 2014 | 97 (28) | 6.4 |

The hypothesis that cases were randomly sampled from the complete network (Table 1: Scenario 1) was inconsistent with the empirical TRAX network (Figure 2A: Scenario 1). Models under this scenario with a high mean degree could reproduce the median of the empirical degree distribution (for mean degree 20, the median was 2.0; Table 3). However, none of these models reproduced the right tail of the empirical degree distribution: at mean degree of 20, the maximum degree in the modeled network was only 20.7, compared to 62 in the empirical network (Table 3). A mean degree greater than 20 in the complete network was required to reproduce the highly connected cases in our transmission study (Table 3; Supplemental Figure 1).

**Figure 2.**
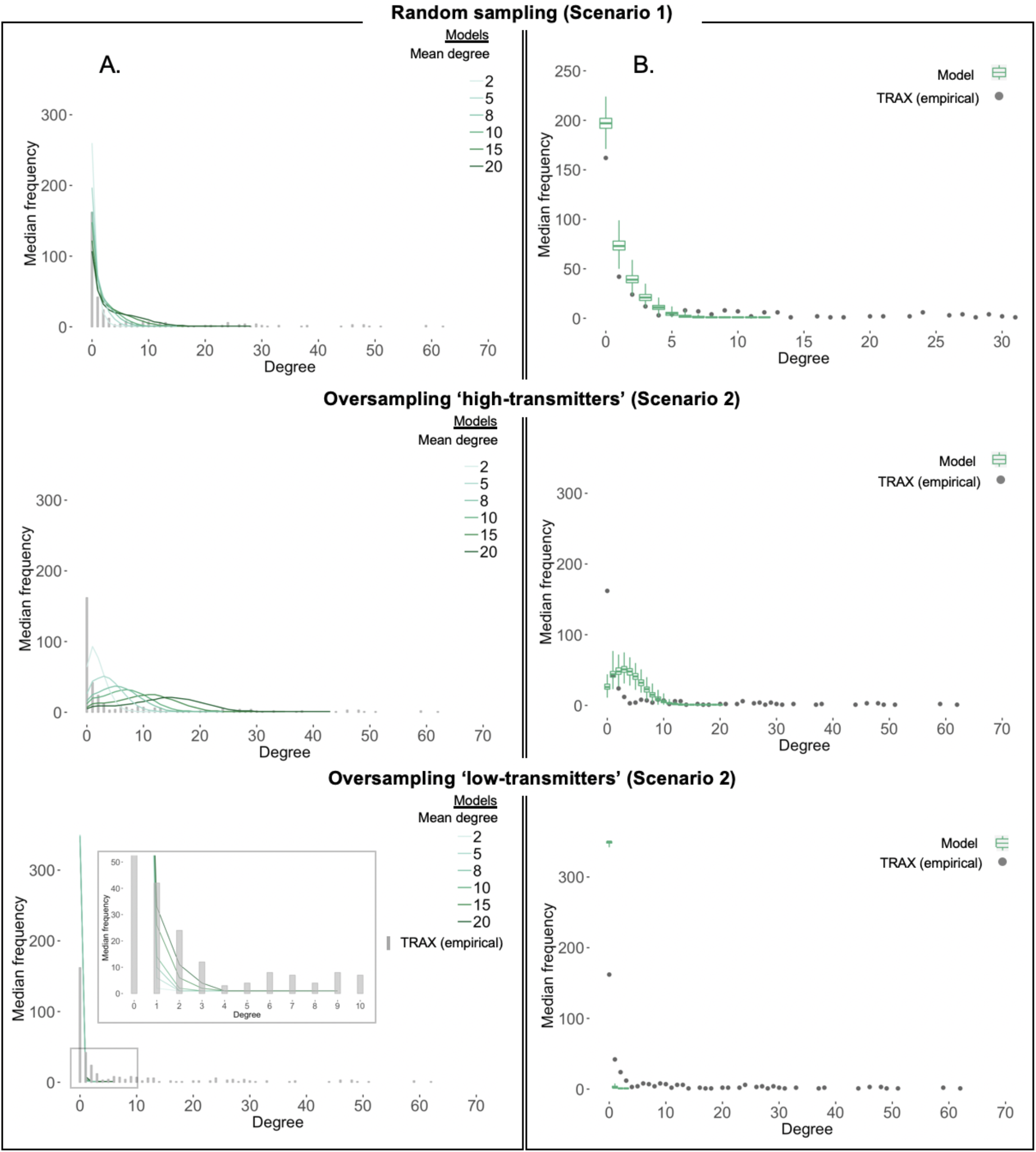
Degree Distributions of Modeled, Sampled Networks Under Scenarios (1) and (2) Degree distributions of empirical (≤ 5 SNPs) and modeled, sampled networks under scenarios 1 and 2. **A.** Grey bars show the distribution of the number of links per case, or the degree distribution, of the empirical network (≤ 5 SNPs) from the TRAX transmission study. Each colored line shows the median degree distribution across 1000 modeled, sampled networks for the corresponding model. Line color indicates the mean degree, or the average number of transmissions per case, assumed in the complete, modeled network. **B**: Range of the degree distributions of the modeled, sampled networks for one model (mean degree = 10). Grey dots show the degree distribution of the empirical network (≤ 5 SNPs) from the TRAX transmission study and are equivalent to the distribution shown by the grey bars in panel A. Colored boxplots show the median, interquartile range, minimum, and maximum frequencies for each degree in the distribution across 1000 modeled, sampled networks.

**Table 3.** Target Statistics for Modeled, Sampled Networks1 Under Scenarios (1) and (2)

|  | Degree, 10th percentile of | Degree, 25th percentile of | Degree, **median** of degree | Degree, 75th percentile of | Degree, **maximum** of |
| --- | --- | --- | --- | --- | --- |
| Mean degree | degree distribution | degree distribution | distribution | degree distribution | degree distribution |
| **TARGET, empirical network (5 SNP)** | **0** | **0** | **1** | **7** | **62** |
| **TARGET, empirical network (3 SNP)** | **0** | **0** | **0** | **1** | **21** |
| **Random sampling (Scenario 1)** |  |  |  |  |  |
| 2 | 0.00 | 0.00 | 0.00 | 0.00 | 4.34 |
| 5 | 0.00 | 0.00 | 0.00 | 1.14 | 7.11 |
| 10 | 0.00 | 0.00 | 1.00 | 2.93 | 11.11 |
| 20 | 0.00 | 0.00 | 2.00 | 6.00 | 20.72 |
| **Oversampling high-transmitters (Scenario 2)** |  |  |  |  |  |
| 2 | 0.00 | 1.00 | 1.93 | 3.00 | 8.78 |
| 5 | 0.95 | 1.97 | 3.65 | 5.61 | 13.59 |
| 10 | 1.72 | 3.97 | 6.87 | 9.76 | 20.09 |
| 20 | 4.01 | 8.85 | 13.85 | 18.16 | 25.97 |
| **Oversampling low-transmitters (Scenario 2)** |  |  |  |  |  |
| 2 | 0.00 | 0.00 | 0.00 | 0.00 | 0.31 |
| 5 | 0.00 | 0.00 | 0.00 | 0.00 | 0.49 |
| 10 | 0.00 | 0.00 | 0.00 | 0.00 | 0.86 |
| 20 | 0.00 | 0.00 | 0.00 | 0.00 | 1.86 |

<sup>1</sup> 1,000 networks were simulated from each model, each modeled network was sampled once.

Scenarios oversampling high- or low-transmitting cases (Scenario 2) significantly changed the structure of modeled networks but did not produce networks fully consistent with the empirical network (Figure 2A: Scenarios 2-3). If high-transmitters were oversampled, the degree distributions of modeled networks were shifted to the right relative to the empirical network (Figure 2A). The 25 ^th^ percentile (range: 1.0–8.9), median (range: 1.9 – 13.9), and 75^th^ percentile (range: 3.0 – 18.2) were close to that of the empirical network (0, 1, and 7, respectively). However, the maximum degree in modeled networks (range: 8.8 – 26.0) could not reproduce the highly-connected cases in the empirical network (degree: 62) (Table 3). When we assumed that low-transmitters were oversampled, the degree distribution was shifted left relative to the empirical network (Figure 2). The median of all modeled networks was at 0 and the maximum degree was much lower (range: 0.3 to 1.9) than the empirical network (Table 3). However, the overall shape of the degree distribution was similar to that of the empirical network, with its mode at 0.

Sampling cases differentially by smear status (Scenario 3) yielded few changes in the degree distributions of modeled networks (Table 4, Figure 3). Results from these models were similar to those from Scenario 1 and did not produce networks with a right tail similar to the empirical network.

**Figure 3.**
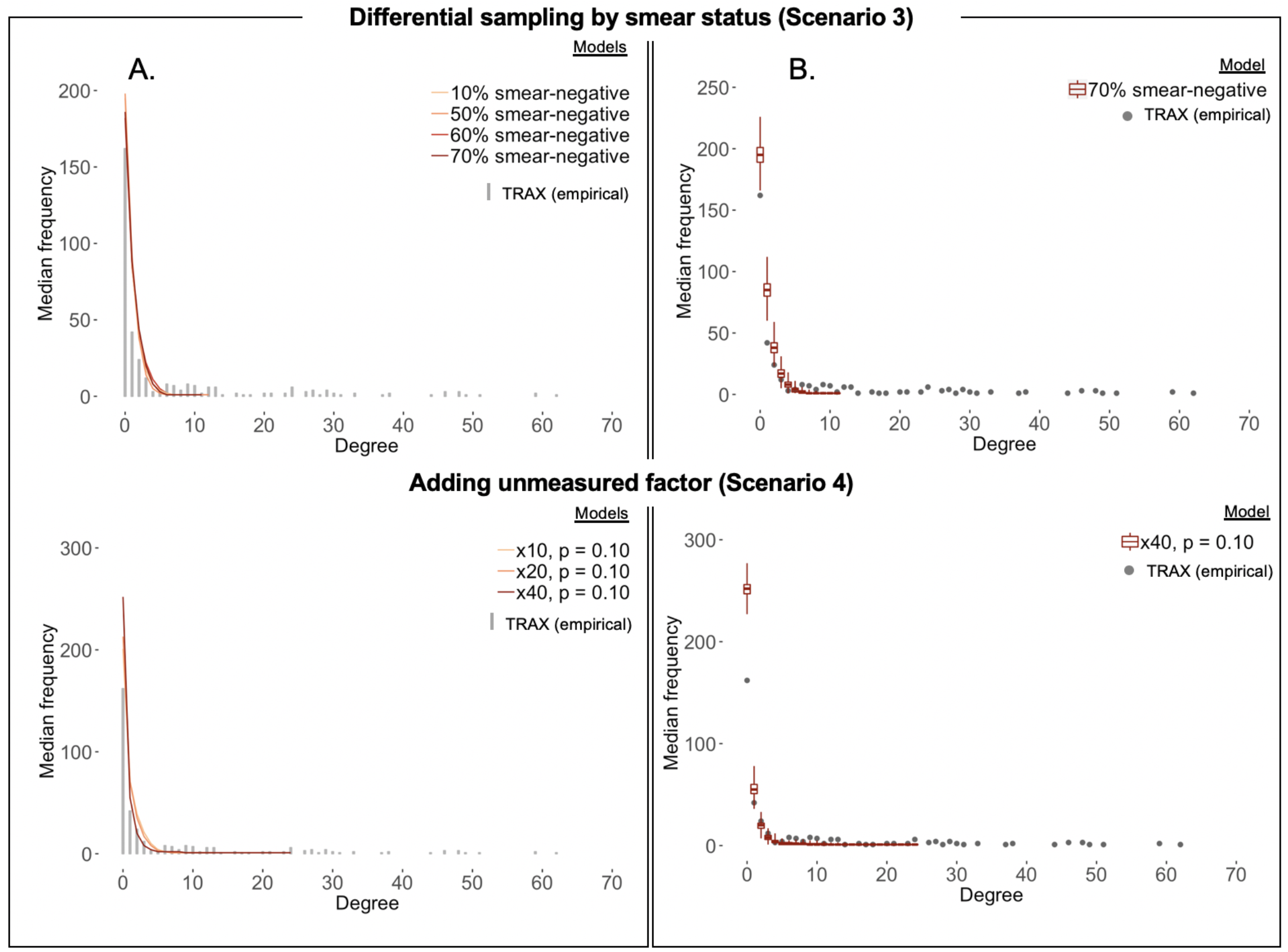
Degree Distributions of Modeled, Sampled Networks Under Scenarios (3) and (4) Degree distributions of empirical (≤ 5 SNPs) and modeled, sampled networks under scenarios 3 and 4. All models shown assume an average mean degree in the complete network of 10. **A.** Grey bars show the distribution of the number of links per case, or the degree distribution, of the empirical network (≤ 5 SNPs) from the TRAX transmission study. Each colored line shows the median degree distribution across 1000 modeled, sampled networks for the corresponding model. Line color indicates the distribution of smear status (Scenario 3) or the strength and prevalence of the unmeasured factor (Scenario 4) assumed in the complete, modeled network. **B**: Range of the degree distributions of the modeled, sampled networks for an individual model. Grey dots show the degree distribution of the empirical network (≤ 5 SNPs) from the TRAX transmission study and are equivalent to the distribution shown by the grey bars inpanel A. Colored boxplots show the median, interquartile range, minimum, and maximum frequencies for each degree in the distribution across 1000 modeled, sampled networks.

**Table 4.** Target statistics of modeled, sampled networks^1^ compared to empirical network under scenarios (3) and (4)

| Mean degree | Degree, 10th percentile<br>of degree distribution | Degree, 25th percentile<br>of degree distribution | Degree, **median** of<br>degree distribution | Degree, 75th percentile<br>of degree distribution | Degree, **maximum** of<br>degree distribution |
| --- | --- | --- | --- | --- | --- |
| **TARGET, empirical network, (5 SNP)** | **0** | **0** | **1** | **7** | **62** |
| **TARGET, empirical network (3 SNP)** | **0** | **0** | **0** | **1** | **22** |
| **Cases sampled preferentially by smear status (Scenario 3)<sup>2</sup>** |  |  |  |  |  |
| 50/50 smear - / + (empirical network: 30/70 smear - / +) |  |  |  |  |  |
| 2 | 0.00 | 0.00 | 0.00 | 0.84 | 4.33 |
| 5 | 0.00 | 0.00 | 0.21 | 1.33 | 7.38 |
| 10 | 0.00 | 0.00 | 1.00 | 2.95 | 10.95 |
| 20 | 0.00 | 0.79 | 2.48 | 5.18 | 15.56 |
| 70/30 smear - / + (empirical network: 30/70 smear - / +) |  |  |  |  |  |
| 2 | 0.00 | 0.00 | 0.00 | 0.44 | 4.12 |
| 5 | 0.00 | 0.00 | 0.02 | 1.01 | 6.84 |
| 10 | 0.00 | 0.00 | 1.00 | 2.41 | 10.83 |
| 20 | 0.00 | 0.72 | 2.17 | 5.04 | 19.06 |
| **Unmeasured factor (Scenario 4)** |  |  |  |  |  |
| 40x, p = 0.10 |  |  |  |  |  |
| 2 | 0.00 | 0.00 | 0.00 | 0.00 | 6.82 |
| 5 | 0.00 | 0.00 | 0.00 | 0.93 | 13.74 |
| 8 | 0.00 | 0.00 | 0.00 | 1.00 | 29.94 |
| 10 | 0.00 | 0.00 | 0.00 | 1.06 | 36.01 |
| 20 | 0.00 | 0.00 | 0.00 | 2.29 | 61.77 |

<sup>1</sup> 1,000 networks were simulated from each model, each modeled network was sampled once.
<sup>2</sup> Smear distribution among TRAX cases (in empirical network): 32% smear-negative, 68% smear-positive.

In the scenario including a latent factor representing superspreading that increased transmission risk 40-fold in 10% of cases (Scenario 4), we could reproduce the target statistics for the maximum of the degree distribution (modeled networks range: 6.8–61.8 vs. empirical network: 62) (Table 4, Figure 3). However, this approach could not reproduce the full empirical degree distribution, in which the 75^th^ percentile was higher (degree: 7) than the modeled networks (range: 0–2.3). (Table 4).

When we assumed larger complete transmission networks (n=4000), modeled networks had a lower median degree (range: 0–1.0) and maximum degree (range: 2.7–9.8) than our primary analysis (n = 2000) (Figure 4, Supplemental Table 2). Smaller complete transmission networks (n=1500) resulted in modeled networks that more closely resembled the empirical network, with a higher median degree (range: 0–2.9) and more highly linked cases (75^th^ percentile: 1.0–8.2, maximum: 5.3–23.4). Thus, the empirical network was more consistent with assumptions of fewer XDR TB cases, rather than more, in the complete network.

**Figure 4.**
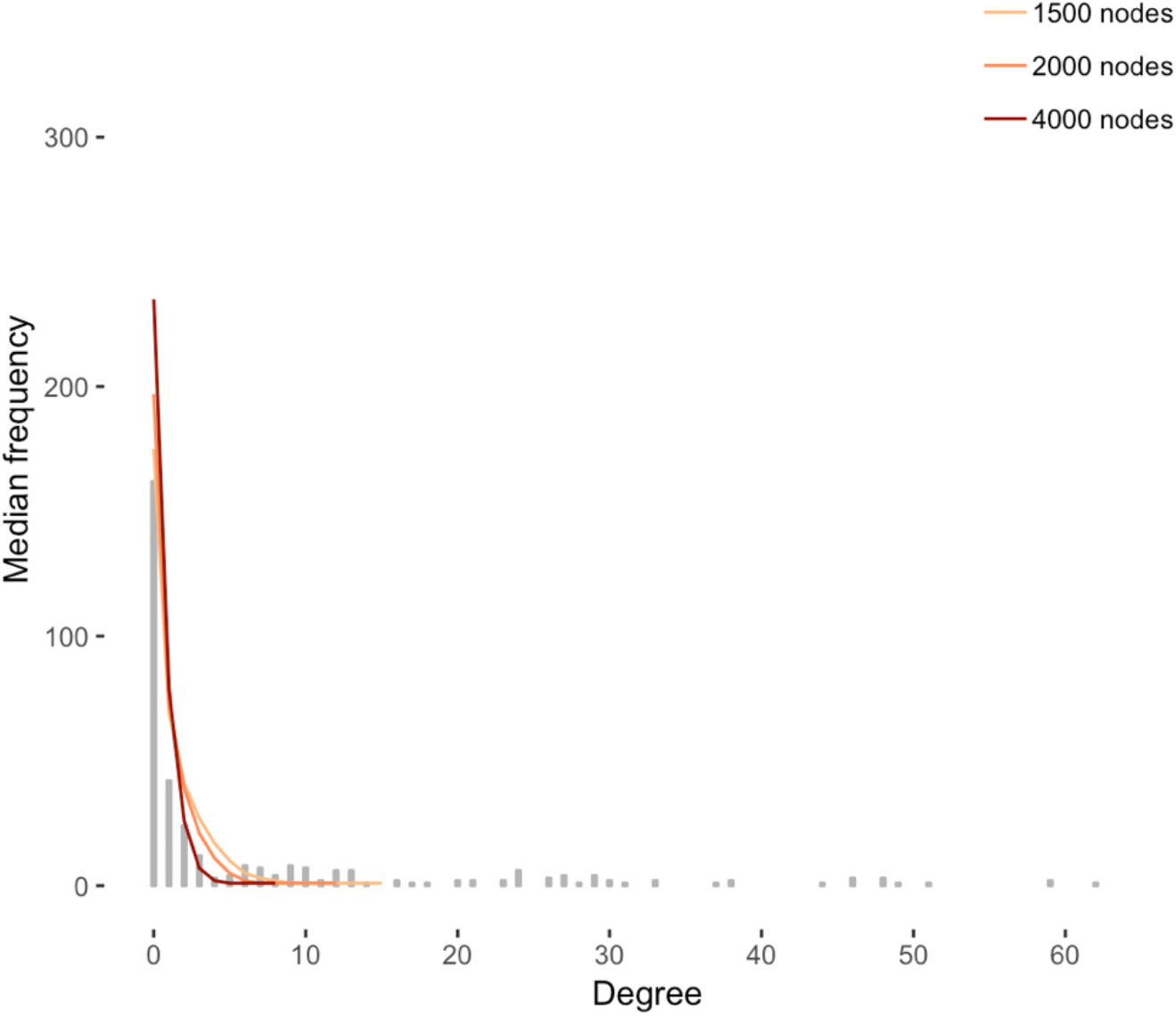
Effect of Modifying Complete Network Size on Modeled, Sampled Networks under Random Sampling. Degree distributions of empirical (≤ 5 SNPs) and modeled, sampled networks under different scenarios. Grey bars show the distribution of the number of links per case, or the degree distribution, of the empirical network (≤ 5 SNPs) from the TRAX transmission study; colored lines show the median degree distribution across 1000 modeled, randomly sampled networks for the corresponding model. Each model makes a different assumption about the total number of XDR TB cases involved in the transmission network during the time period of our transmission study (2011-2014), or the size of the complete transmission network. The model shown has a mean degree in the complete network of 10.

Finally, we assessed the robustness of our results to the SNP threshold used to define a genomic edge. Since a threshold of 3 SNPs requires cases’ TB strains to be more closely related to define an edge, the empirical degree distribution is shifted to the left relative to that of the network defined by a 5 SNP threshold (Figure 5). Under random sampling, the model with a mean degree of 20 could almost reproduce the maximum (20.7) of the empirical 3 SNP network (22 in the empirical network), but not the median (2 in the modeled network, 0 in the empirical network) (Table 3).

**Figure 5.**
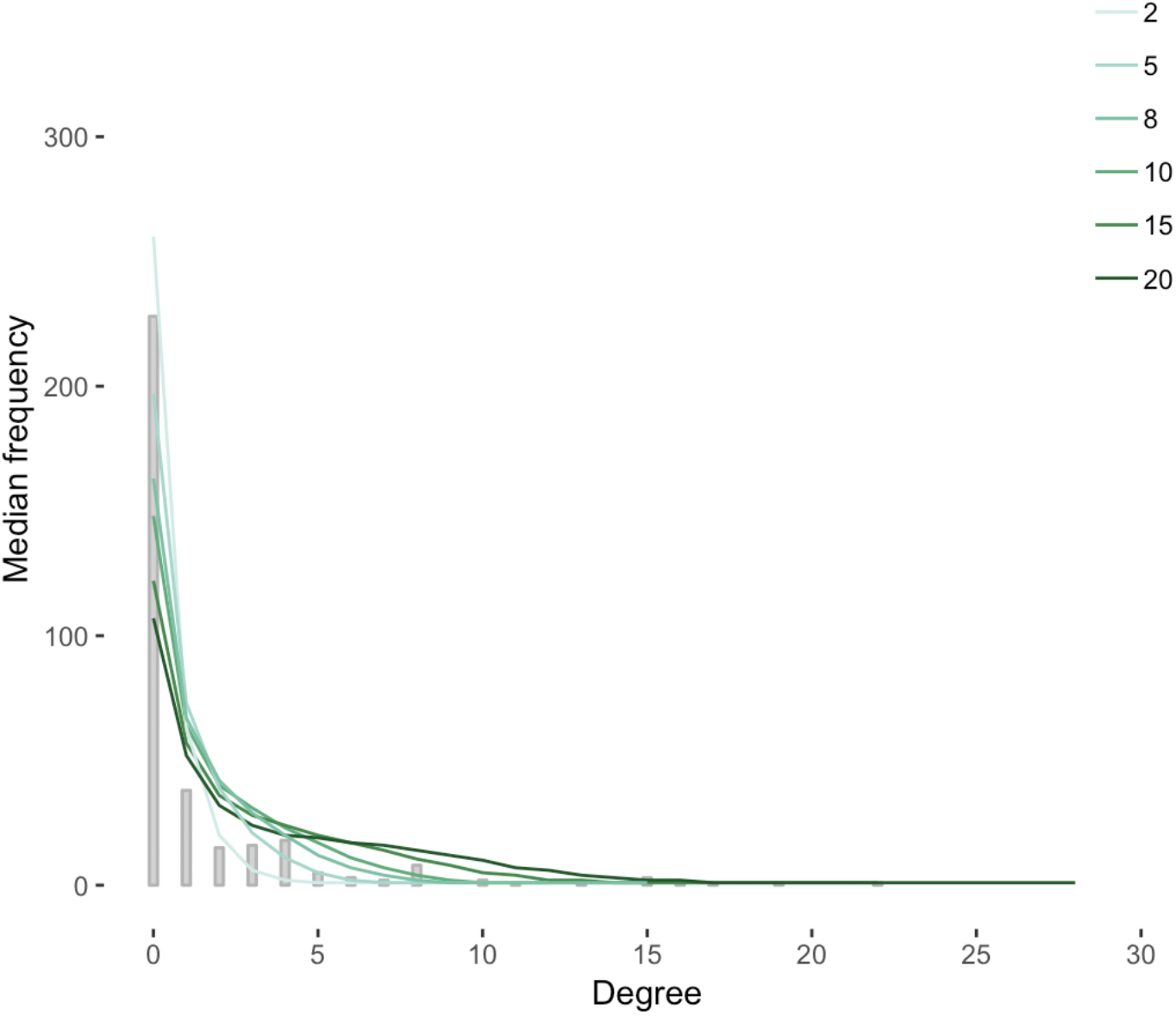
Effect of Reducing SNP Threshold (≤ 3 SNPs) on the Empirical Network. Degree distributions of empirical (≤ 3 SNPs) and modeled, sampled networks under scenarios 1 and 2. Grey bars show the distribution of the number of links per case, or the degree distribution, of the empirical network (≤ 3 SNPs) from the TRAX transmission study. Each colored line shows the median degree distribution across 1000 modeled, randomly sampled networks for the corresponding model. Line color indicates the mean degree, or the average number of transmissions per case, assumed in the complete, modeled network.

When we accounted for a ‘superspreading’ factor (Scenario 4), modeled networks could indeed reproduce a degree distribution similar to the 3 SNP empirical network. At a mean degree of 8, the modeled network matched closely with the empirical network, with the median at 0 (empirical network: 0), 75^th^ percentile at 1.0 (empirical network: 1), and maximum at 29.9 (empirical network: 22) (Table 4).

## DISCUSSION

In this study, we explored the extent to which partial and non-random sampling of TB transmission events may bias inferences we aim to make from transmission networks constructed using incomplete molecular and epidemiologic data. We determined that missingness in our transmission study was unlikely to be random; rather, we likely oversampled low-transmitting cases. We also found evidence that superspreading behavior may partially account for the structure of the transmission network we observed. The methodological framework outlined this study sheds light on the key assumptions required to make inference from incomplete sampling of TB cases. Our results advise caution when interpreting transmission networks measured from incomplete data in TB-endemic settings without a clear understanding of the sampling frame and factors potentially contributing to bias.

That cases were missing completely at random from our transmission study was unlikely based on our models, and this finding was generally robust to our choice of SNP threshold to define transmission. Instead, we found that low-transmitting cases were more likely to be sampled than high-transmitting cases. There are several potential explanations for this finding: first, there is sufficient Second, specific behaviors that lead to higher contact rates, including the common practice of traveling to urban centers for seasonal employment, may also lead to a lower likelihood of diagnosis and retention in care.(25–28)

It is striking that even with systematic oversampling of *low*-transmitting cases, we found evidence of superspreading, or cases that were responsible for many transmission events. The characteristics of TB ‘superspreaders’ are not well-understood. Superspreading may be caused by social or behavioral factors leading to many transmission events, such as cross-province travel for employment or frequent use of public transport during one’s infectious period.(19, 29) Other host factors may include yet unmeasurable clinical or biological features: for example, recent evidence suggests that certain individuals are more likely to generate infectious, *Mtb*-containing aerosols independent of their clinical disease presentation.(30, 31) Alternatively, superspreaders may be infected with an exceptionally transmissible variant.(32, 33) Future research should focus on identifying drivers of superspreading to inform interventions targeting specific locations or activities linked with superspreading events.

Our results were sensitive to factors about which there is substantial uncertainty in TB, including key natural history features and the SNP threshold defining a direct transmission event. Our primary models varied the mean degree in the complete network from 2 to 20. This range was selected after considering previous estimates of the reproduction number of TB (R_0_), which is not well-characterized.(34) Interestingly, the models most consistent with the empirical network had a mean degree of 10 and above, which is substantially higher than most previous estimates of R_0_. This suggests either that R_0_ is truly higher in this setting or that our definition of a transmission event, at 5 SNPs, may be too lenient.(9) To address this issue, we examined networks defined using a 3 SNP threshold. This empirical network was consistent with a wider range of tested models than the network based on a 5 SNP threshold. This result emphasizes the challenges of relying upon pairwise genomic distances to define transmission events: conclusions regarding transmission can be vastly different based on the threshold being used.

Lastly, our results were sensitive to assumptions about the total number of XDR TB cases comprising the complete network. Underdiagnosis of TB is a persistent challenge in low-resource settings and is even more difficult for XDR TB, which requires culture-based drug susceptibility testing. In our primary analysis, we assumed that approximately half of all XDR TB cases are diagnosed. We found that larger complete networks were less likely to match the empirical network, suggesting that it is unlikely we greatly underestimated the number of XDR TB cases in KwaZulu-Natal. However, the results from this sensitivity analysis underscore the broader challenge of understanding the true magnitude of TB disease burden in low-resource settings and using this information to accurately model population-level transmission dynamics.

### Limitations

First, we did not distinguish the direction of transmission in modeled or empirical networks in order to avoid fitting of our models with uncertain parameter data. More sophisticated probabilistic methods to define genomic transmission links between cases that account for directionality may be warranted in future analyses.(35) Second, we did not explicitly account for the fact that XDR TB cases may have acquired resistance (i.e., been diagnosed with XDR TB after failing treatment for a less resistant infection). These cases would have contributed to forward XDR TB transmission but would not have a source case with XDR TB. While this is an important feature of drug-resistant TB epidemiology, the vast majority of XDR TB cases are due to transmission rather than acquired resistance.(6, 7) Lastly, ERGMs utilize Poisson distributions (conditional on nodal attributes and other network features) to model the number of edges per node; this distribution may fail to capture inherent properties of TB transmission. Recent studies have shown that the distribution of R_0_ of TB may be best represented by a negative binomial distribution, which may more accurately capture the effect of superspreading on the network degree distribution.(17) While ERGMs allow for model building that accounts for multiple interacting generative mechanisms for edge formation, our future work will investigate alternative network model specifications with relaxed distributional assumptions.

### Conclusions

While a clearer understanding of transmission is critical in settings with a high burden of disease, sparse data in these settings poses serious challenges for interpretation of transmission studies. This is, to our knowledge, the first study to use network modeling approaches to understand the impact of missingness in a study of TB transmission. Although the scenarios we tested did not exactly reproduce the transmission network observed in our study, we found evidence that superspreading behavior and biased sampling may explain the observed network. Future research should focus on identifying the specific host, pathogen, or environmental factors that contribute to superspreading. Further, future transmission studies in high-incidence settings should aim to understand the impact of incomplete and potentially biased sampling more explicitly and identify key assumptions regarding missingness on which inferences are based. These efforts will allow more accurate mapping of TB transmission patterns in endemic settings, where the need to design effective interventions tailored to local epidemics is greatest.

## Supporting information

Technical Appendix

