## Supplementary material for "Modeling missing cases and transmission links in networks of extensively drug-resistant tuberculosis in KwaZulu-Natal, South Africa": Technical Appendix

Supplemental Technical Appendix, Nelson et al.


**1 Introduction**

*1.1 Model framework*

*1.2 Model software*

**2 Empirical Data**

**3 Defining models using missing case assumptions**

*3.1 Cases missing at random*

*3.2 Cases missing by level of connectivity*

*3.3 Cases missing by HIV/smear status*

*3.4 Unmeasured factor contributing to transmission*

**4 Clinical measures**

*5.1 Cough duration*

*5.2 Smear status*

*5.3 HIV*

*5.4* Mtb *strain type*

**5 Demographic measures**

*6.1 Age*

**6 Joint distributions**

*7.1 Age and smear status*

*7.2 Age and HIV*

*7.3 Other (Smear status and HIV)*

**7 Simulation and sampling methods**

**8 Sensitivity analyses**

*9.1 Genomic (SNP) threshold for transmission*

*9.2 Full network size*

**9 References**

### 1 Introduction

This technical appendix describes the models used in the associated manuscript, including their conceptual basis and parameterization as well as simulation procedures and statistical analysis.

#### *1.1 Model framework*

The network models in this study were used to represent and simulate transmission networks of active tuberculosis (TB) cases. Links, or edges, in modeled networks represent a transmission event that occurred between two cases in the network.

Modeled networks do not involve individuals (1) *infected* with TB but whom did not progress to active disease or (2) exposed contacts of TB cases. Rather, modeled networks reflect all transmission events observed from a sample of cases enrolled in our transmission study of XDR TB, which enrolled patients over a four-year period. Note that this network does not include transmission events that occurred either before the study period began or after it ended.

Moreover, this network is comprised of cases with extensively drug-resistant (XDR) TB. Importantly, not all cases of XDR TB were infected with XDR TB; rather, they acquired resistance as a result of inadequate treatment of a more drug-susceptible strain of TB. For simplicity, we ignore this feature of drug-resistant TB epidemiology when modeling the transmission networks in this study. Moreover, it has been shown that the vast majority of XDR TB are due to transmission of already-resistant strains, rather than acquired resistance.

In the model, each case in the network was assigned specific clinical and demographic attributes according to a pre-defined distributions based on the data collected in our transmission study. (For example, we specified the age distribution of cases in the complete network based on the known age distribution of TB cases in our transmission study.) The attributes assigned to each case influence the number of other cases to whom that case is connected in the network. (See the Empirical Data section for more detail.)

We included each attribute as a 'nodefactor' term in the network model. (Some nodefactor terms represented the joint distributions of two attributes, see Joint Distributions section.) This allowed the number of links to vary by an individual's attributes. We defined 'target statistics' for the number of links a given case had in the network based on their demographic and clinical attributes. To do this, we calculated the number of edges in the network corresponding to each nodefactor term, or the number of edges in the network involving nodes with that particular attribute. (For example, the target statistic for the HIV nodefactor term was the number of edges involving HIV-positive nodes, or the number of transmission events in the network involving HIV-positive cases.)

For attributes with well-studied effects on infectiousness (i.e., smear status), we used estimated from the literature to define target statistics for the corresponding nodefactor term in our models. For attributes specific to our study or this population (i.e., time of enrollment in study or strain type) we used empirical data from the TRAX network to calculated target statistics for nodefactor terms (see Empirical Data section below).

#### *1.2 Model software*

All models were programmed in R. The code used to create and analyze these models is available on GitHub. (https://github.com/kbratnelson/tb-ergms)

The modeling methods employed in this study utilized the *ergm* R package, which requires the *statnet* suite of software.

#### 2 Empirical Data

The Transmission of XDR TB (TRAX) study is a cross-sectional study that enrolled 404 XDR TB cases from KwaZulu-Natal province, South Africa from 2011-2014. This study collected clinical, demographic and social network data from enrolled cases. The primary aim of this study was to estimate the proportion of XDR TB cases due to transmission, as compared to those due to acquired resistance. The major finding of this study was that at least 70% of XDR cases in this settings are due to transmission.[1]

The empirical transmission network was created from 344 cases with available whole genome sequencing results of their *Mtb* isolates. We created sequencing-based networks using pairwise differences between *Mtb* sequences. We considered fewer than 5 single nucleotide polymorphisms a transmission link and constructed an undirected network in which each node represented a case and each edge represented a transmission link based on a pairwise SNP difference threshold. We considered several SNP thresholds, as the appropriate threshold to define a direct transmission event between two cases may vary by setting (see Sensitivity Analyses section below).

In this empirical undirected network, the maximum degree was 62, there were 162 (47%) of cases with no links (degree = 0), and 62 (18%) of cases with 10 or more links (degree ≥ 10). See Table 1 in the manuscript and Supplemental Table 1 for more descriptive characteristics of the empirical network defined by 5 and 3 SNPs, respectively.

To define the likelihood of being linked in the modeled networks based on a particular attribute, we used estimates from the literature of relative infectiousness by clinical or demographic characteristics. Using this information, we defined the relative mean degree of cases with each attribute as compared to a reference group. (For example, we defined the mean degree of HIV-positive relative to HIV-negative cases based on estimate from the literature on the relative infectiousness of HIV-positive as compared to HIV-negative cases.)

To calculate the target statistics for each nodefactor term in the network model, we multiplied the relative mean degree for each attribute by the overall mean degree of the network. To scale this according to the number of nodes in the network to this attribute, we then multiplied that value by the proportion of nodes in that network with the attribute. (For example, to model a network with an overall mean degree of 5, we multiplied the mean degree of HIV-positive cases by 5, then by the proportion of HIV-positive cases in the network, to give the total number of edges associated with HIV-positive cases in the network.) Models with target statistics specified for all levels of every attribute did not easily converge, so we reduced the number of target statistics for attributes with more than four categories. For these variables, we used the 3–4 categories corresponding to the highest number of edges as target statistics to parameterize models. Using these target statistics, we simulated full transmission networks from each model.

#### 3 Defining models using missing case assumptions

We defined models and simulated full transmission networks under scenarios which made different assumptions about cases missing from the empirical TRAX network.

*3.1 Cases missing at random*

We assumed that cases missing at random would result in missing transmission links randomly across the network. Therefore, to simulate full transmission networks based on the assumption cases were missing at random, we used the same model but modified the mean degree in the full network. We simulated full networks with mean degrees of 2, 5, 8, 10, 15, and 20, 50, 100, and 200. Results from models of networks with mean degree of 2 through 20 are presented in the main manuscript and models with mean degree greater than 20 are presented in the supplemental results.

The degree of a given case in the network is the sum of the links to that case, (in theory, one, representing the source case), and all forward transmission links from that case. Since only cases of active disease are represented in the network, the mean degree therefore roughly corresponds to the mean number of secondary cases caused by a TB case in the network (less one, corresponding to the source case of their infection) *and progressed to disease during the study period*.

It is important to note that the modeled and simulated networks are undirected, so the direction of transmission is not indicated. Parameterizing models to include both risk factors for infection and transmission was beyond the scope of this project.

*3.2 Cases missing by level of connectivity*

We assumed two opposing scenarios: that cases who were highly connected in the full network were more likely to be sampled, and that cases that were poorly connected in the full network were more likely to be sampled. To simulate the former scenario, we created and used sampling weights proportional to each cases' degree in the full network; to simulate the latter scenario, we created and used sampling weights inversely proportional to degree in the full network.

We sampled cases using this method in full networks with various mean degrees (2, 5, 8, 10, 15, 20).

*3.3 Cases missing by HIV/smear status*

We assumed that cases were undersampled, or oversampled, systematically based on their smear status. We modified the distribution of smear status in the full network relative to the empirical TRAX network to reflect each hypothesis. In the TRAX study, 70% of XDR TB cases were HIV-positive. We created scenarios in which the proportion of HIV-positive cases in the full network was 10%, 40%, and 90%. A smaller proportion of cases in the full network relative to the empiric TRAX network (10%, 40%) reflects the assumption that HIV-positive cases were oversampled; a larger proportion (90%) reflects the assumption HIV-positive cases were undersampled.

We sampled cases from full networks with varying distributions of HIV and smear status with various mean degrees (2, 5, 8, 10, 15, 20).

*3.4 Unmeasured factor contributing to transmission*

We hypothesized that a factor contributing strongly to transmission risk but that was not accounted for in our model (a 'superspreading' factor) might have a substantial impact on network structure. We hypothesized that such a factor might increase transmission by at least 10 times, that is, cases with this factor would be responsible for 10 times as many transmission events as those lacking this factor. We created a 'nodefactor' term in the model for this unmeasured, superspreading factor at various strengths (10x, 20x, 40x) and varied its prevalence in the population of cases from 10 to 30%. The results of models assuming the strongest effect of the factor (x40) in 10% of cases are presented in the main manuscript.

#### 4 Size of full networks

To simulate full networks, it was necessary to make assumptions about the number of cases involved in XDR TB transmission over the time period 2011-2014. We estimated the number of diagnosed and undiagnosed XDR TB cases in KwaZulu-Natal province contributing to transmission using data from the South African National Tuberculosis Drug Resistance Survey.[2] We then used active case-finding studies to estimate the proportion of TB cases in South Africa that are undiagnosed.[3]

*332, 783 TB cases in SA in 2014*
*Proportion of cases with pulmonary TB (infectious form) = 0.89*
*Proportion of cases in KwaZulu-Natal province (area of study) = 0.31*
*Proportion of cases with XDR = 0.005*

*332, 783 * (0.31) * (0.89) * (0.005) * 4 yrs = 1836 cases (736 - 2572)*

Accounting for underdiagnosis of TB cases [3], multiply by factor of 2:

*1836 cases * 2 = 3672 cases (1472 - 5144)*

For our primary analysis, we estimated a total number of XDR TB cases on the lower end of this range. We simulated networks assuming a full network size of n = 2000, but also explored the impact of changing network size (see Sensitivity Analyses section below).

#### 5 Clinical measures

##### *5.1 Cough duration*

We categorized cough duration by month. The distribution of cough duration and target statistics for the mean degree in each group are below:

| Cough duration | n (%) | Mean degree (Source: TRAX) |
|---|---|---|
| No cough* | 128 (37) | 5.0 |
| 1 month* | 60 (17) | 6.5 |
| 2 months* | 51 (15) | 8.1 |

| | | |
|---|---|---|
| 3 months* | 72 (21) | 8.1 |
| 4 months | 16 (5) | 4.6 |
| 5 months | 17 (5) | 3.7 |

* Target statistics defined for these categories

As there was little information on the effect of cough duration on transmission in the literature, we used the mean degree from the empirical TRAX network to define target statistics for modeled networks. We used target statistics for the largest categories 'No cough', '1 month', '2 months', and '3 months' in network models.

*5.2 Smear status*

Although both smear status and grade were available, we used only smear status (smear-positive and smear-negative) to reduce the number of model parameters. The marginal distribution is below:

| Smear status | n (%) | Mean degree (Source: Abu-Raddad et al.[4]) |
|---|---|---|
| Negative | 109 (32) | 1 |
| Positive | 235 (68) | 4 |

The mean degree parameters are based on relative infectiousness estimates by Abu-Raddad et al. that we normalized, assuming a mean degree of 1 in the smear-negative group. We used the joint distribution of age and smear status for model target statistics; see Joint Distributions section.

*5.3 HIV*

Although both HIV status and information on virologic suppression were available, we used only HIV status (HIV-positive and HIV-negative) to reduce the number of model parameters. The marginal distribution is below:

| HIV status | n (%) | Mean degree |
|---|---|---|
| Negative | 78 (23) | 1 |
| Positive | 266 (77) | 1 |

Since there is conflicting evidence as to whether HIV-positive or HIV-negative individuals are more infectious, we chose to assume no difference in infectiousness, and therefore the mean degree for HIV-positive and HIV-negative individuals was the same. We used the joint distribution of age and HIV status for model target statistics; see Joint Distributions section.

*5.4* Mtb *strain type*

The dominant strain of XDR TB in KwaZulu-Natal is the LAM4 strain. There is evidence that the phenotype of this strain may lead to differences in its transmission and evolutionary rate. [5,6] We categorized *Mtb* strains into LAM4 or non-LAM4.

| *Mtb* strain | n (%) | Mean degree (Source: TRAX) |
|---|---|---|
| LAM4* | 259 (23) | 8.3 |
| Non-LAM4 | 85 (77) | 0.2 |

* Target statistics defined for these categories

Since there was little evidence in the literature about the relative infectiousness of LAM4 and non-LAM4 strains of XDR TB, we used the relative mean degrees estimated from the empirical TRAX network for modeled networks.

#### 6 Demographic measures

*6.1 Age*

We categorized age into four groups: 0-15, 16-34, 35-54, >55.

| Age category | n (%) | Mean degree (Source: Wood et al.[7]) |
|---|---|---|
| 0 - 15 | 12 (3) | 1 |
| 16 – 34 | 171 (50) | 1.58 |
| 35 – 54 | 134 (39) | 0.98 |
| > 55 | 27 (8) | 0.75 |

The mean degree parameters are based on relative infectiousness estimates by Wood et al. that we normalized, assuming a mean degree of 1 in the 0-15 age group. We used the joint distribution of age/HIV status and age/smear status for model target statistics; see Joint Distributions section.

#### 7 Joint distributions

*7.1 Age and smear status*

| Age category | Smear status | n (%) | Mean degree |
|---|---|---|---|
| 0 - 15 | Negative | 7 (2) | 1.00 |
| 16 – 34* | Negative | 44 (13) | 1.58 |
| 35 – 54* | Negative | 40 (12) | 0.98 |
| > 55 | Negative | 18 (5) | 0.75 |
| 0 - 15 | Positive | 5 (1) | 4.00 |
| 16 – 34* | Positive | 107 (31) | 6.32 |
| 35 – 54* | Positive | 79 (23) | 3.92 |
| > 55 | Positive | 7 (2) | 3.00 |

* Target statistics defined for these categories

We calculated the mean degree by multiplying the relative infectiousness measures for smear status and age group from the Tables in Section 5.2 and 6.1, respectively. (We assumed independence of the two measure of infectiousness.) We used target statistics for the largest categories, ‘16-34, smear-negative’, ‘35-54, smear-negative’, ‘16-34, smear-positive’, and ‘35-54, smear-positive’ as target statistics for network models.

*7.2 Age and HIV*

| Age category | HIV status | n (%) | Mean degree |
|---|---|---|---|
| 0 - 15 | Negative | 5 (1) | 1.00 |
| 16 - 34 | Negative | 41 (12) | 1.58 |
| 35 - 54 | Negative | 15 (14) | 0.98 |
| > 55 | Negative | 17 (5) | 0.75 |
| 0 - 15 | Positive | 7 (2) | 1.00 |
| *16 - 34 | Positive | 130 (38) | 1.58 |
| *35 - 54 | Positive | 119 (35) | 0.98 |
| > 55 | Positive | 10 (3) | 0.75 |

* Target statistics defined for these categories

We calculated the mean degree by multiplying the relative infectiousness measures for HIV status and age group from the Tables in Section 5.3 and 6.1, respectively. (We assumed independence of the two measure of infectiousness.) We used target statistics for the largest categories, '16-34, HIV-positive', and '35-54, HIV-positive' as target statistics for network models.

*7.3 Other (Smear status and HIV)*

Although smear-negative disease tends to be more common among HIV-positive TB cases, we did not find this association in the empirical data. The proportion of cases with HIV was nearly equivalent among smear-positive and smear-negative cases and the proportion of smear-positive cases was nearly equivalent among HIV-positive and HIV-negative cases. Thus, we chose not to represent the joint distribution of smear status and HIV in model target statistics.

#### 8 Simulation and sampling methods

From each network model, we simulated 1000 networks. We specified the following parameters of the Markov Chain Monte Carlo (MCMC) algorithm: we set the number of burn-in simulations as 100000, the MCMC interval as 5000, and the MCMC sample size as 10000.

We ensured that the MCMC algorithm used to estimate parameters for each model converged appropriately by checking for adequate mixing of the MCMC chain and sufficient exploration of parameter space using the mcmc.diagnostics function in the *ergm* package.

We sampled 350 cases from each simulated, full network, mimicking sampling 350 cases in our TRAX study from the larger population of XDR TB cases. We compared the degree distributions of modeled, sampled networks to that of the empirical TRAX network.

We attempted to 'match' the following quantiles of the empirical degree distribution: (1) $10^{th}$ percentile; (2) $25^{th}$ percentile; (3) median ($50^{th}$ percentile), (4) $75^{th}$ percentile, and (5) maximum ($100^{th}$ percentile).

#### 9 Sensitivity analyses

*9.1 Genomic threshold for transmission*

Since the threshold for defining genomic evidence of transmission is not well-defined, we also defined an empirical network using a more stringent threshold of 3 pairwise SNP differences. This resulted in no changes to modeled networks, but did change the target statistics we attempted to 'match' with modeled, sampled networks. The differences in the empirical networks defined by different SNP thresholds can be examined by comparing Table 1 in the main manuscript (5 SNP threshold) and Supplemental Table 1 (3 SNP threshold). The target statistics for both networks are shown in Tables 3 and 4.

*9.2 Full network size*

Given the uncertainty around the total number of XDR TB cases contributing to transmission, we considered several other sizes of the full network. We assumed that the full network may be larger than 2000 cases (n = 4000 cases), or that it may be smaller (n = 1500 cases). We compared the results from these networks to our main models, which assumed a full network size of 2000 cases.
