## Supplementary Material for "Modeling missing cases and transmission links in networks of extensively drug-resistant tuberculosis in KwaZulu-Natal, South Africa"

**Supplemental Table 1**. Descriptive characteristics, sequencing-based network of XDR TB cases in the TRAX study (≤3 SNP threshold)

| | N (%) | Mean |
|---|---|---|
| Total network | | |
| Edges (genomic links) | 240 | - |
| Isolates (unlinked cases) | 228 (66) | - |
| Overall mean degree | - | 1.4 |
| 10th percentile | 0 | |
| Median degree (IQR) | 0 (0,1) | |
| Maximum degree | 22 | |
| Nodes with degree ≥ 10 | 9 (3) | |
| By attribute | | |
| **HIV status** | | |
| HIV- | 78 (23) | 1.15 |
| HIV+, undetectable VL | 133 (39) | 1.50 |
| HIV+, detectable VL | 133 (39) | 1.37 |
| **Cough duration** | | |
| No cough | 128 (37) | 1.30 |
| 1mo | 60 (17) | 1.35 |
| 2mo | 51 (15) | 1.75 |
| 3mo | 72 (21) | 1.69 |
| 4mo | 16 (5) | 0.06 |
| 5mo | 17 (5) | 0.71 |
| **Smear status/grade** | | |
| Negative | 109 (32) | 1.49 |
| Scanty + | 37 (11) | 1.73 |
| Positive, grade 1 | 59 (17) | 1.05 |
| Positive, grade 2 | 51 (15) | 1.35 |
| Positive, grade 3+ | 88 (26) | 1.31 |
| **Sex** | | |
| Female | 202 (59) | 1.34 |

| | | |
|---|---|---|
| Male | 142 (41) | 1.42 |
| **Age category** | | |
| < 15 | 12 (3) | 1.25 |
| 16-34 | 171 (50) | 1.19 |
| 35-54 | 134 (39) | 1.53 |
| > 55 | 27 (8) | 1.78 |
| **TB Strain** | | |
| HP | 259 (75) | 1.79 |
| Other | 85 (25) | 0.09 |
| **Year** | | |
| 2011 | 58 (17) | 1.84 |
| 2012 | 107 (31) | 1.36 |
| 2013 | 82 (24) | 1.10 |
| 2014 | 97 (28) | 1.34 |

**Supplemental Figure 1**. Mean degree required to reproduce the maximum degree in the empirical network

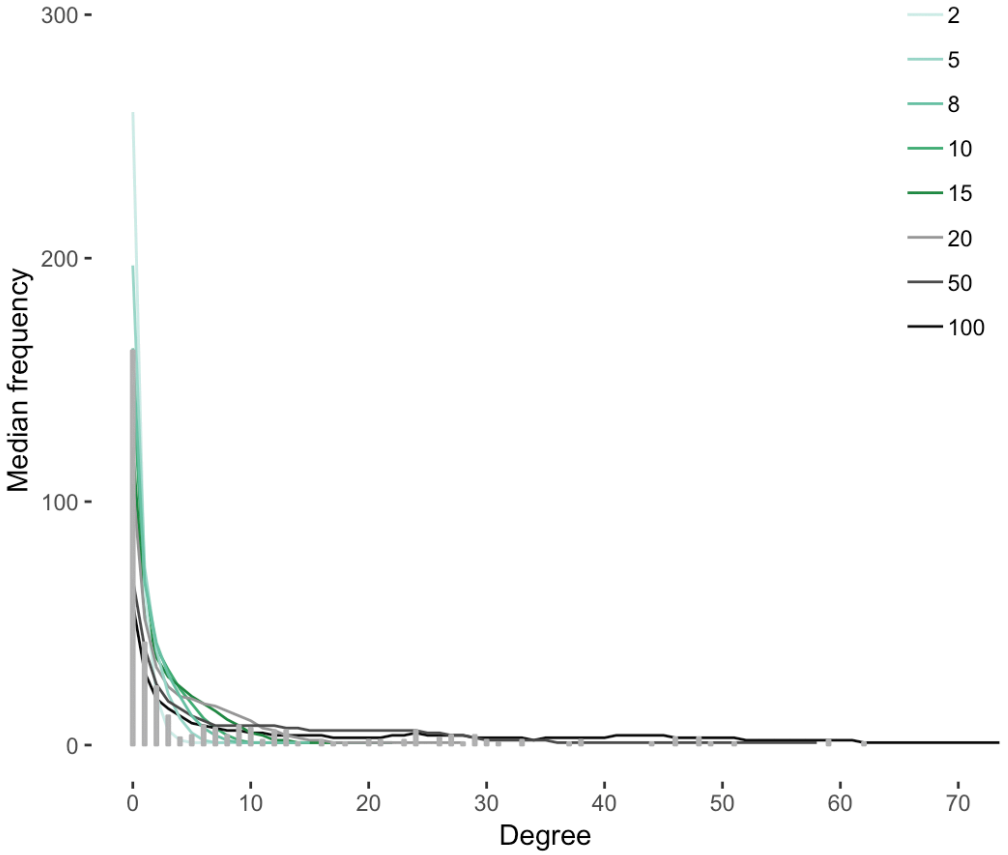

**Supplemental Figure 1**. Mean degree required to reproduce the maximum degree in the empirical network. Grey bars show the distribution of the number of links per case, or the degree distribution, of the empirical network (≤ 5 SNPs) from the TRAX transmission study. Each colored line shows the median degree distribution across 1000 modeled, sampled networks for the corresponding model. Line color indicates the mean degree, or the average number of transmissions per case, assumed in the complete, simulated network.

**Supplemental Table 2**. Effect of modifying network size on modeled, sampled networks

| Mean degree | Degree, 10th percentile of degree distribution | Degree, 25th percentile of degree distribution | Degree, 50th percentile (**median**) of degree distribution | Degree, 75th percentile of degree distribution | Degree, 100th percentile (**maximum**) of degree distribution |
|---|---|---|---|---|---|
| **TARGET (5 SNP)** | **0.00** | **0.00** | **1.00** | **7.00** | **62.00** |
| **Random sampling (Scenario 1) with network size = 4000** | | | | | |
| 2 | 0.00 | 0.00 | 0.00 | 0.00 | 2.71 |
| 5 | 0.00 | 0.00 | 0.00 | 1.00 | 4.26 |
| 8 | 0.00 | 0.00 | 0.01 | 1.00 | 5.46 |
| 10 | 0.00 | 0.00 | 0.49 | 1.50 | 6.43 |
| 15 | 0.00 | 0.00 | 1.00 | 2.13 | 7.94 |
| 20 | 0.00 | 0.00 | 1.04 | 3.02 | 9.75 |
| **Random sampling (Scenario 1) with network size = 2000 (From Table 3)** | | | | | |
| 2 | 0.00 | 0.00 | 0.00 | 0.00 | 4.34 |
| 5 | 0.00 | 0.00 | 0.00 | 1.14 | 7.11 |
| 8 | 0.00 | 0.00 | 0.94 | 2.06 | 9.57 |
| 10 | 0.00 | 0.00 | 1.00 | 2.93 | 11.11 |
| 15 | 0.00 | 0.00 | 1.22 | 4.21 | 14.91 |
| 20 | 0.00 | 0.00 | 2.00 | 6.00 | 20.72 |
| **Random sampling (Scenario 1) with network size = 1500** | | | | | |
| 2 | 0.00 | 0.00 | 0.00 | 1.00 | 5.27 |
| 5 | 0.00 | 0.00 | 0.49 | 1.99 | 9.16 |
| 8 | 0.00 | 0.00 | 1.00 | 3.00 | 12.41 |
| 10 | 0.00 | 0.00 | 1.01 | 3.94 | 14.44 |
| 15 | 0.00 | 0.00 | 2.00 | 5.98 | 18.91 |
| 20 | 0.00 | 0.12 | 2.90 | 8.17 | 23.43 |

[1] 1,000 networks were simulated from each model, each simulated network was sampled once.
